# Phylogenomic relationships and historical biogeography in the South American vegetable ivory palms (Phytelepheae)

**DOI:** 10.1101/2020.09.03.280941

**Authors:** Sebastián Escobar, Andrew J. Helmstetter, Rommel Montúfar, Thomas L. P. Couvreur, Henrik Balslev

**Author notes:** Thomas L. P. Couvreur and Henrik Balslev should be considered joint senior author. **Correspondence**, Sebastián Escobar, Section for Ecoinformatics and Biodiversity, Department of Biology, Aarhus University, DK 8000 Aarhus C, Denmark.

## Abstract

The vegetable ivory palms (Phytelepheae) form a small group of Neotropical palms whose phylogenetic relationships are not fully understood. Three genera and eight species are currently recognized; however, it has been suggested that *Phytelephas macrocarpa* could include the species *Phytelephas seemannii* and *Phytelephas schottii* because of supposed phylogenetic relatedness and similar morphology. We inferred their phylogenetic relationships and divergence time estimates using the 32 most clock-like loci of a custom palm bait-kit formed by 176 genes and four fossils for time calibration. We additionally explored the historical biogeography of the tribe under the recovered phylogenetic relationships. Our fossil-dated tree showed the eight species previously recognized, and that *P. macrocarpa* is not closely related to *P. seemanii* and *P. schottii*, which, as a consequence, should not be included in *P. macrocarpa*. The ancestor of the vegetable ivory palms was widely-distributed in the Chocó, the inter-Andean valley of the Magdalena River, and the Amazonia during the Miocene at 19.25 Ma. Early diversification in *Phytelephas* at 5.27 Ma can be attributed to trans-Andean vicariance between the Chocó/Magdalena and the Amazonia. Our results support the role of Andean uplift in the early diversification of *Phytelephas* under new phylogenetic relationships inferred from genomic data.

## 1. Introduction

The vegetable ivory palms (Phytelepheae) are a clade of eight South American lowland or premontane rain forest species. Phytelepheae species are distributed in northwestern South America and adjacent Panama. Most species are restricted to either side of the Andes within the Chocó, the inter-Andean valley of the Magdalena River (hereinafter referred as Magdalena), or the Amazonia (Supporting Information Figure S1). These are single stemmed dioecious palms with pinnate leaves (Barfod, 1991). They produce large (~5 cm in length), white, and extremely hard seeds that are harvested and commercialized because of their resemblance to animal ivory (Barfod, 1991). The vegetable ivory palms were considered a distant and highly-evolved group of palms because of their “aberrant” floral morphology. Family wide molecular phylogenetic studies have, however, shown that the Phytelepheae clustered within the subfamily Ceroxyloideae (Asmussen et al., 2006; Baker et al., 2009; Dransfield et al., 2008; Trénel et al., 2007). Inference of phylogenetic relationships based on few genetic markers (3 to 5) supported the existence of three genera within Phytelepheae, the monospecific *Ammandra* and *Aphandra*, and *Phytelephas* with six species (Barfod et al., 2010; Trénel et al., 2007).

The latest molecular phylogenetic tree of Phytelephae was based on nrDNA ITS2, PRK, and RPB2 nuclear sequences (Barfod et al., 2010). The authors dated the phylogenetic tree using fossil, geological and secondary calibrations points for Ceroxyloideae (Trénel et al., 2007). They inferred that the genus *Phytelephas* diverged from the sister genera *Ammandra* and *Aphandra* during the Miocene between 17–25 Ma (Trénel et al., 2007). *Phytelephas aequatorialis* Spruce and *Phytelephas tumacana* O.F. Cook have been inferred as sister species, similarly to *Phytelephas seemanii* O.F. Cook and *Phytelephas schottii* H. Wendl., which in turn form a sister clade to *Phytelephas macrocarpa* Ruiz & Pav. (Barfod et al., 2010; Trénel et al., 2007). It has been suggested that *P. seemanii* and *P. schottii* are intraspecific entities of *P. macrocarpa* because of morphological similarities (Bernal and Galeano, 2010), implying the existence of only four species within *Phytelephas*. The position of *Phytelephas tenuicaulis* (Barfod) A.J. Hend. is inconsistent between both phylogenetic analyses, either being sister to *P. aequatorialis* and *P. tumacana* (Barfod et al., 2010), or sister to *P. macrocarpa, P. seemanii* and *P. schottii* (Trénel et al., 2007). This phylogenetic disagreement in turn results in different biogeographic inference for Phytelepheae.

Based on ancestral area reconstructions, the ancestral range of Phytelepheae during the Miocene was inferred to be the tropical rain forests of the Chocó, Magdalena, and Amazonia (Barfod et al., 2010). At that time the western Andes reached less than half of their current height and were suggested not to be a barrier between trans-Andean rain forests in the Chocó and the Amazonia (Gregory-Wodzicki, 2000; Hoorn et al., 2010). The Chocó and Magdalena rain forests were already separated by the western Andes during the Miocene; however, both regions remained virtually connected in the lowlands of northwestern Colombia (Hoorn et al., 2010). The western Andes uplifted rapidly at ~10 Ma isolating the Chocó from the Amazonia, whereas the eastern Andes in Colombia started to uplift at ~5 Ma separating Magdalena from Amazonia (Gregory-Wodzicki, 2000). Early diversification in *Phytelephas* is not explained by trans-Andean vicariance but instead by adaptive shifts of morphological traits (Barfod et al., 2010). Posterior diversification in *Phytelephas* can be explained by three events of Andean vicariance: two events between the Chocó and Amazonia, and one more event between the Chocó and Magdalena. Under an alternate scenario (Trénel et al., 2007), trans-Andean vicariance occurred also late in *Phytelephas* between the Chocó and the Amazonia first and then between the Chocó and Magdalena. These two different scenarios of evolutionary history in *Phytelephas* arise from non-similar phylogenetic topologies. Thus, it seems necessary to infer more realistic phylogenetic relationships in *Phytelephas* to understand the influence that the Andes have had in their diversification.

Here, we infer the phylogenetic relationships and divergence time estimates of the vegetable ivory palms using genomic data and two individuals per species. Additionally, we estimated their ancestral range and historical biogeography. We aim to test if *P. macrocarpa, P. seemanii*, and *P. schottii* are inferred as a well-supported clade, which could provide support for the inclusion of these entities under the name *P. macrocarpa*. We also test the impact that Andean uplift has had in the diversification of the vegetable ivory palms in northwestern South America and adjacent Panama.

## 2. Materials and methods

### 2.1 Sampling and DNA extraction

We sampled two individuals of each of the eight species of tribe Phytelepheae (Table 1). We used leaves dried in silica gel when possible, otherwise we extracted DNA from herbarium dried leaves available at AAU Herbarium (Aarhus University). We sampled one individual of *Ammandra decasperma* from each side of the Andes (Chocó and Amazonia) to explore the intraspecific trans-Andean diversification of this species. We did the same for *Phytelephas macrocarpa*, which is hypothesized to represents a trans-Andean distribution (Bernal and Galeano, 2010). We sampled 19 additional species from the palm subfamilies Coryphoideae, Arecoideae, and Ceroxyloideae as outgroup. We isolated total genomic DNA using the MATAB and chloroform separation methods (Mariac et al., 2014). We then checked DNA quality and quantity with a NanoDrop™ One spectrophotometer (Thermo Scientific^™^).

**Table 1.**
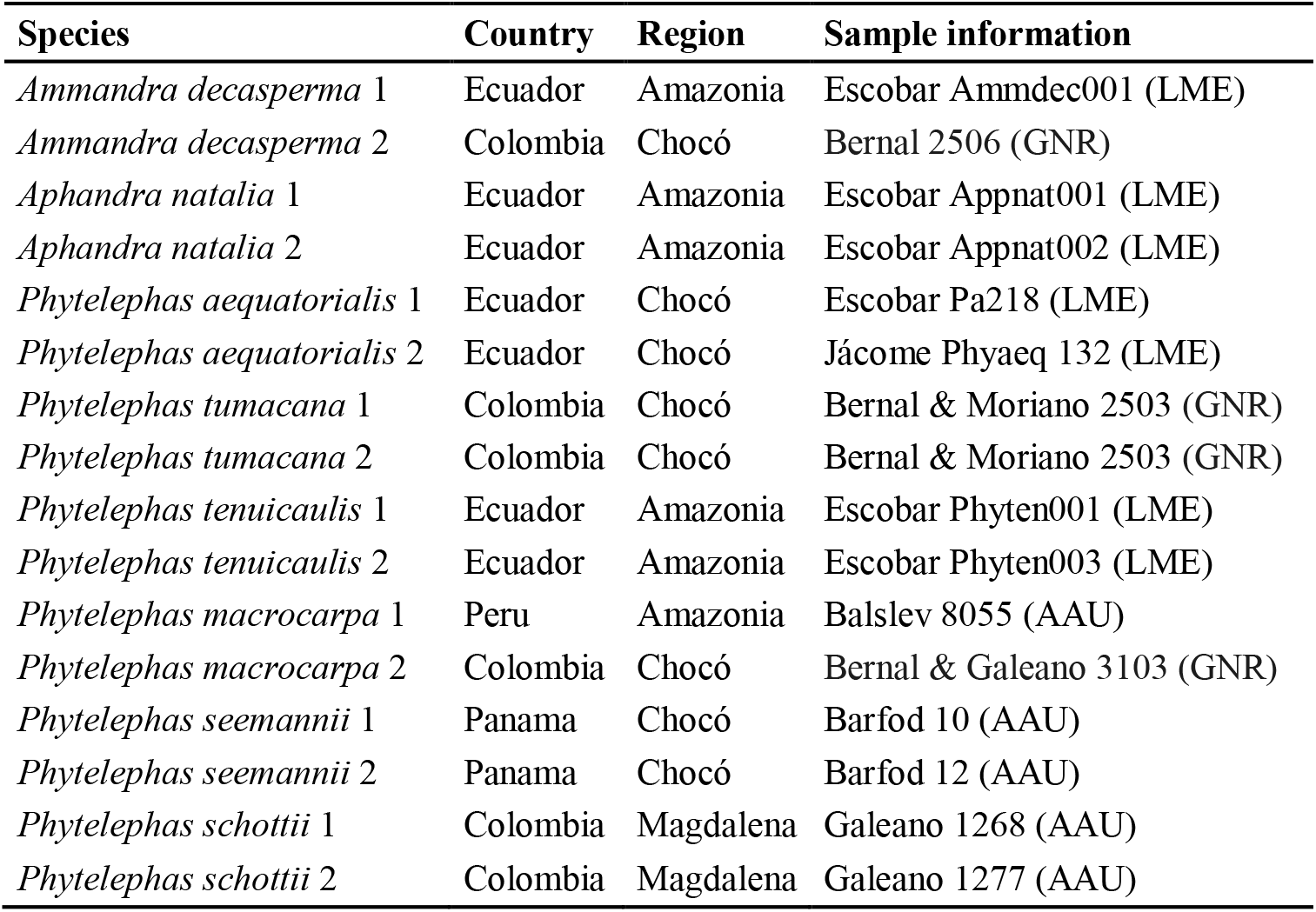
List of individuals used to construct a fossil-dated phylogenetic tree of the vegetable ivory palms (Phytelepheae). LME = Lab of Molecular Ecology, Herbarium QCA, PUCE, Ecuador; GNR = Guadalito Natural Reserve, Colombia; AAU = AAU Herbarium, Aarhus University, Denmark.

### 2.2 Sequencing and data processing

We performed laboratory work at the facilities of the *Institut de Recherche pour le Développement* (IRD) in Montpellier, France. Genomic libraries were prepared following Rohland and Reich (2012) and used a palm custom bait kit to capture 176 nuclear genes (Heyduk et al., 2016). Total DNA was sheared to a mean target size of 350 bp with a Bioruptor^®^ Pico sonication device (Diagenode). DNA were repaired, ligated, and nick filled-in before being amplified for 10 cycles as a prehibridization step. We cleaned and quantified the DNA before bulking the samples in two libraries. We added biotin-labeled baits to the libraries to hybridize the targeted regions (Heyduk et al., 2016). Streptavidin-coated magnetic beads were used to immobilize the hybridized biotin-labeled baits. We amplified the enriched DNA fragments in a RT-PCR during 15 cycles to complete adapters. Libraries were sequenced with an Illumina^®^ MiSeq (paired-end, length 150 bp) at the CIRAD (Montpellier, France).

The South Green Platform at the IRD was used for high-performance computing analyses. We controlled the quality of the reads using fastQC (http://www.bioinformatics.babraham.ac.uk/projects/fastqc). We used a 0-mismatch threshold for demultiplexing using the script *demultadapt* (https://github.com/Maillol/demultadapt) and removed the adapters using cutadapt v.1.2.1 (Martin, 2011). We discarded the reads with length < 35 bp and quality mean values (Q) < 30 using a custom script (https://github.com/SouthGreenPlatform/arcad-hts/blob/master/scripts/arcad_hts_2_Filter_Fastq_On_Mean_Quality.pl). We paired *forward* and *reverse* sequences using an adapted script (Tranchant-Dubreuil et al., 2018). Quality was checked one more time using fastQC. The *fastx* script (https://github.com/agordon/fastx_toolkit) was used to remove the last 6 bp of the *reverse* sequences to ensure the removal of barcodes in sequences < 150 bp.

We used Hybpiper v.1.2 (Johnson et al., 2016) for processing the data. We identified paralogous loci by building exon trees of potential paralogs flagged by Hybpiper in RAxML v.8.2.9 (Stamatakis, 2014) using a GTR+G model, as in Helmstetter et al. (2020). If > 50% of exons belonging to a certain gene were determined as paralogs, then the gene was marked as “true” paralog and removed from further steps. We kept only the loci with > 75% of their length reconstructed in > 75% of the individuals. The remaining supercontigs were aligned using the “--auto” option in MAFFT v.7.305 (Katoh and Standley, 2013) and were cleaned using Gblocks v.0.91b (Castresana, 2000). New gene trees were inferred in RAxML using a GTR+G model, which were rooted using the most distant species to *P. aequatorialis (Phoenix rupicola, Latania loddigesii*, and *Hyphaene thebaica*) in PHYX (Brown et al., 2017). We then calculated the tip-to-root variance using the function *get_var_length* from SortaDate (https://github.com/FePhyFoFum/SortaDate/blob/master/src/get_var_length.py) and chose the 32 most clock-like loci to reduce computational time (Smith et al., 2018). The corresponding 32 *fasta* alignments were converted to *nexus* format using PGDSpider v.2.1.1.5 (Lischer and Excoffier, 2012).

### 2.3 Phylogenomic reconstruction and divergence time estimation

We inferred the phylogenetic relationships and estimated divergence times in the vegetable ivory palms using a fossil calibration approach. The phylogenetic tree was constructed using Bayesian inference in BEAST v.2.5.2 (Bouckaert et al., 2019). We used a relaxed clock log normal prior and set the clock rate to 0.001, making sure that the ‘estimate’ box was checked. We set the best nucleotide substitution model of evolution for each alignment as determined in ModelTest (https://github.com/ddarriba/modeltest). The Yule model for interspecific analyses was used as the tree prior. To estimate the divergence times between the branches of our tree, we used four fossil calibration points previously used in other palm studies (Couvreur, Forest, & Baker, 2011; Trénel et al., 2007), with one placed within Phytelepheae. More information on the fossils and their position can be found in Table 2 and Figure 1. We set uniform distributions for the calibration points because exponential distributions did not reach convergence for all effective sample sizes (ESS). An additional monophyly prior was added to the subfamily Arecoideae because it was not inferred as such during test runs, even though its monophyly has been previously established (Baker et al., 2009; Faurby et al., 2016). We performed three independent Markov Chain Monte Carlo (MCMC) analyses of 300 million generations sampling every 30,000 trees. We checked that the MCMC runs reached convergence using Tracer v.1.7 (Rambaut et al., 2018), ensuring that all ESS > 200. The log and tree files of the three runs were combined with LogCombiner v2.5.2 (Bouckaert et al., 2019) with a burn-in of 10%. TreeAnnotator v.2.5.2 (Bouckaert et al., 2019) was used to generate a maximum clade credibility (MCC) tree by summarizing the remaining 27,003 trees, and to calculate nodes’ ages and 95% highest posterior density intervals (95% HPD) using the option ‘Keep target heights’. The resulting tree was visualized in FigTree v.1.4 (http://tree.bio.ed.ac.uk/software/figtree/).

**Figure 1.**
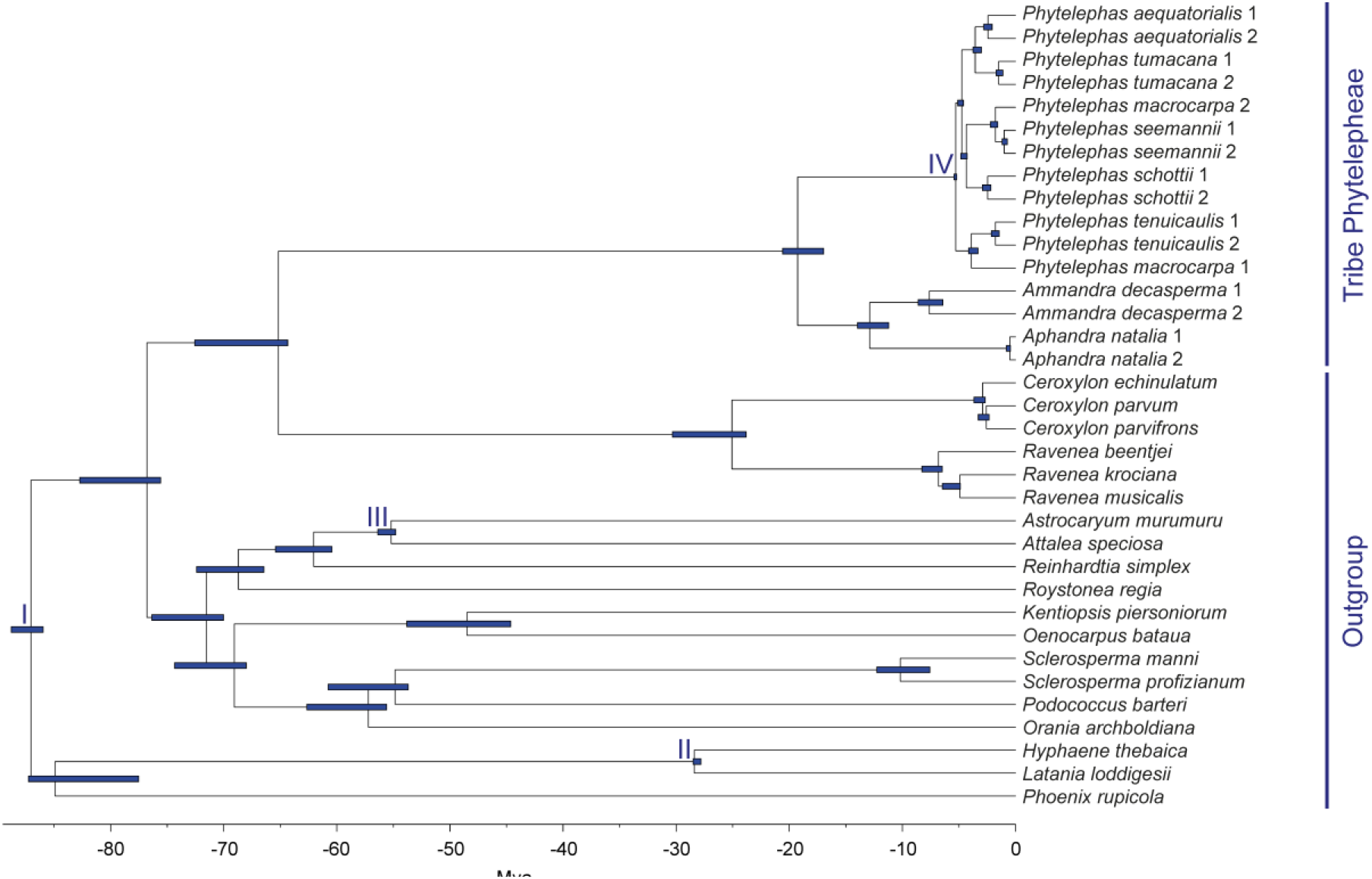
Fossil-dated phylogenetic tree of the vegetable ivory palms (Phytelepheae). The tree was inferred using 32 clock-like genes and Bayesian inference in BEAST. Blue lines represent the 95% higher posterior densities of time estimates. The roman numbers I-IV represent the position of four fossils used as calibration points to date the tree (Table 1).

**Table 2.** Fossils used to date the phylogenetic tree of the vegetable ivory palms (Phytelepheae). Lower and upper bounds were set as uniform.

| Position in the tree | Fossil name | Lower (Ma) | Upper (Ma) | Monophyly prior |
| --- | --- | --- | --- | --- |
| I | *Sabalites carolinensis* | 85.8 | 88.8 | Yes |
| II | *Hyphaene kapelmanii* | 27 | 28.5 | Yes |
| III | Attaleinae | 54.8 | 60.79 | Yes |
| IV | *Phytelephas olsonni* | 5.2 | Infinite | Yes |

### 2.4 Biogeographic inference

We defined biogeographic regions based on the distribution of Phytelepheae species. To visualize their distribution, we compiled occurrence records from the Global Biodiversity Information Facility (GBIF), the Berkeley Ecoengine (Ecoengine), iNaturalist, and the Botanical Information and Ecological Network (BIEN v.3). The R packages *BIEN* (Maitner et al., 2018) and *spocc* (Chamberlain and Boettiger, 2017) were used to access the records. We removed records that were recorded prior to 1950, were duplicated, had coordinate uncertainty greater than 5 km, had impossible or unlikely coordinates, were within 10 km of a country capital or political unit centroids, and were within 100 m of biodiversity institutions. Given the low number of species, we were able to clean the downloaded data manually following specialized literature (Barfod, 1991; Barfod et al., 2010; Bernal & Galeano, 2010; Borchsenius, Borgtoft Pedersen, & Balslev, 1998; Trénel et al., 2007). Additional records were obtained from our own and private databases of colleagues. We defined three biogeographical regions based on the recovered distribution of Phytelepheae (Supporting Information Figure S1): Chocó, Magdalena, and Amazonia.

The biographic history and range evolution of Phytelepheae was inferred using the R package *BioGeoBEARS* 0.2.1 (Matzke, 2013). We first used the package *ape* (Paradis and Schliep, 2019) to keep only one individual of each Phytelepheae species in addition to three *Ceroxylon* species used as outgroups. The three models of range evolution implemented in *BioGeoBEARS*, dispersal-extinction-cladogenesis (DEC), BAYAREALIKE, and DIVALIKE were tested as well as their variants that implement a jump parameter + J. The maximum number of areas that any species can occupy was set to three because *A. decasperma* is present in the three regions (Bernal & Galeano, 2010; Supporting Information Figure S1). We tested a biogeographic model associated with the uplift history of the central and northern Andes. To do so, we constructed dispersal cost matrices between the three defined biogeographic regions under three time periods (Supporting Information Table S1) based on the orogenic history of the Andes (Gregory-Wodzicki, 2000; Hoorn et al., 2010). The first period from 0–5 Ma when the eastern Andes in Colombia uplifted rapidly and impeded dispersal between Magdalena and the Amazonia. Western Andes was a dispersal barrier between the Chocó and the Amazonia at this time. From 5–10 Ma the western Andes reached their actual height and started acting as a dispersal barrier. From 10–70 Ma the western Andes reached half of their current height, allowing trans-Andean dispersal between the Chocó and Amazonia. We defined an additional ‘null’ biogeographic model with equal probability of dispersal between regions through time. Thus, a total of 12 models of range evolution were compared using the Akaike information criterion (AIC) and the AIC corrected for small sample size (AICc). The best model was selected based on the lowest AIC and AICc values.

## 3. Results

### 3.1 Phylogeny and divergence time estimation

We generated a total of 20.37 million reads from the 35 individuals representing 27 palm species. HYBPIPER detected 133 loci with > 75% of their length reconstructed in > 75% of individuals and 16 loci with signs of paralogy, reducing our dataset to 117 supercontigs. All nodes within Phytelepheae showed a posterior support of 1 in the fossil-dated tree (Figure 1). The crown node of Phytelepheae was estimated to 19.25 Ma (95% HPD 16.96–20.58). The genera *Ammandra* and *Aphandra* diverged 12.88 Ma (95% HPD 11.2–13.97), while trans-Andean individuals of *A. decasperma* diverged 7.61 Ma (95% HPD 6.42–8.61). The crown node of the genus *Phytelephas* was estimated at 5.27 Ma (95% HPD 5.2–5.43). All *Phytelephas* individuals grouped with their conspecifics, except *P. macrocarpa*. The individual described as *P. macrocarpa* from Chocó grouped with *P. seemannii*, whereas the individual from Amazonia resolved as sister of *P. tenuicaulis* (Figure 1).

### 3.2 Biogeographic inference

After cleaning the data, we retained and visualized 813 occurrence records of Phytelepheae (Figure 2; Supporting Information Figure S1). The DEC model stratified under three time periods was determined as the best range evolution model based on the AIC and AICc values (Supporting Information Table S2). A unique widespread region formed by the Chocó, Magdalena, and Amazonia was estimated as the ancestral range of Phytelepheae, of the clade formed by *Ammandra* and *Aphandra*, and of the genus *Phytelephas* (Figure 2). Early diversification in *Phytelephas* divided its ancestral range between the Chocó/Magdalena and the Amazonia. The ancestral range of Phytelepheae and *Ceroxylon* was estimated in a region formed by the Amazonia and the Andes; however, the certainty of this estimation was low (Figure 2).

**Figure 2.**
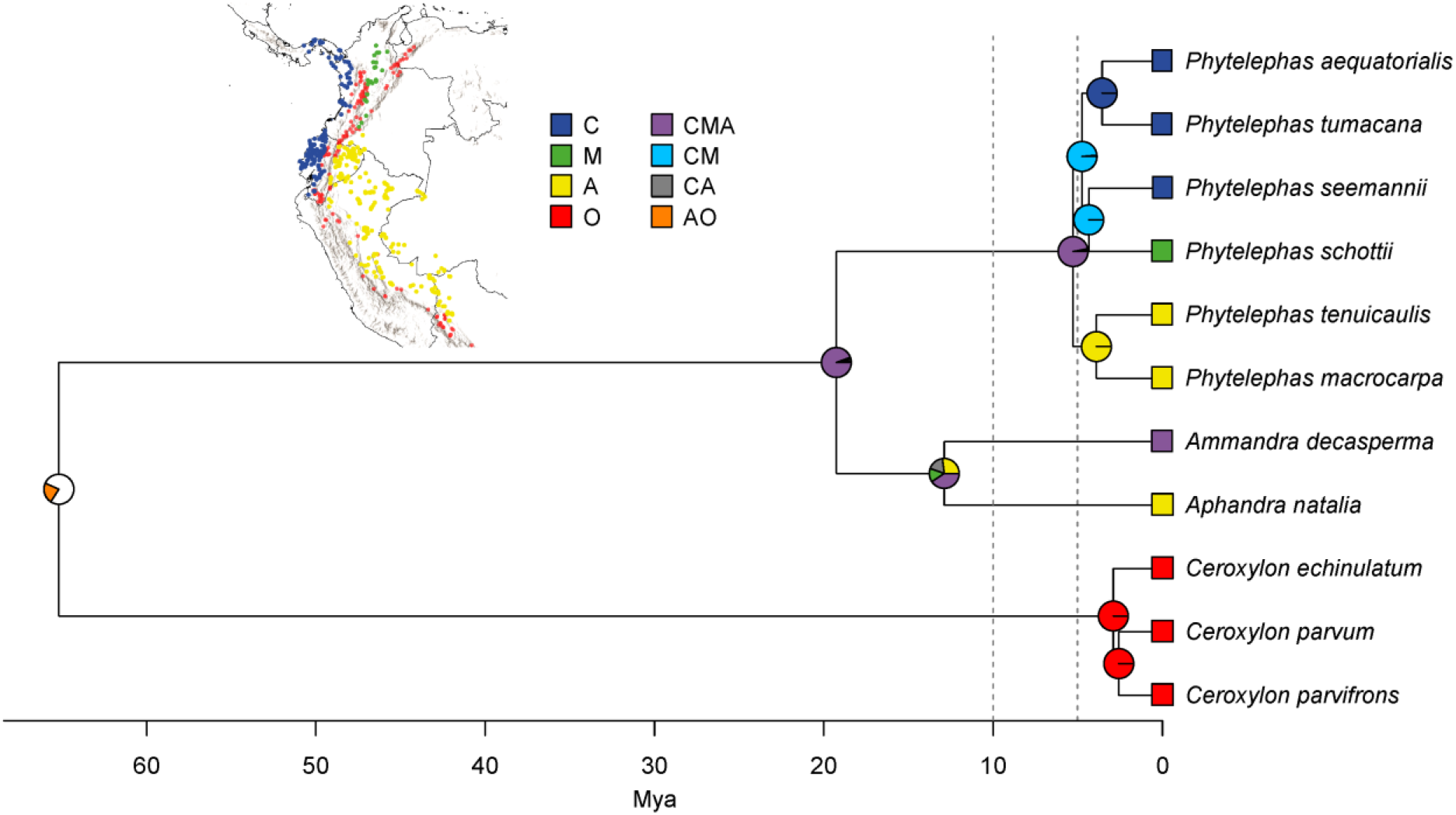
Biogeographic history of the vegetable ivory palms (Phytelepheae) under the dispersal-extinction-cladogenesis (DEC) model of range evolution in BioGeoBears. Inset map shows the distribution of Phytelepheae species (Supporting information Figure S1) in the Chocó (C), the inter-Andean valley of the Magdalena River (M), and the Amazonia (A). It also shows the distribution of the three *Ceroxylon* species in the Andes (O), which were used as outgroup. The analysis was stratified under three time periods based on the uplift history of the Andes (Gregory-Wodzicki, 2000) and was performed using dispersal cost matrices between regions (Supporting information Table S1). Gray dotted lines show the approximate time when the western Andes became a barrier between the Chocó and the Amazonia (~10 Ma) and when the eastern Andes isolated Magdalena from the Amazonia (~5 Ma).

## 4. Discussion

We reconstructed the phylogenetic relationships of the vegetable ivory palms (tribe Phytelepheae) using a phylogenomic approach. Phytelepheae was resolved as monophyletic (Figure 1), similar to previous molecular reconstructions (Barfod et al., 2010; Trénel et al., 2007). Eight previously recognized species of vegetable ivory palms (Barfod et al., 2010; Trénel et al., 2007) can be inferred from our fossil-dated tree (Figure 1), with six species within the genus *Phytelephas*. The position of the two *P. macrocarpa* individuals in the tree (Figure 1) suggests that they do not belong to the same species. The phylogenetic relatedness of the *P. macrocarpa* individual from the Chocó with the individuals described as *P. seemannii* from Panama (Figure 1) suggests that this individual belongs to *P. seemannii*. We did not analyze individuals described as *P. macrocarpa* from Magdalena (Bernal and Galeano, 2010); however, the individuals of *P. schottii* from Magdalena formed a clade separated from the *P. macrocarpa* individual from the Amazonia (Figure 1). Thus, our phylogenetic results do not support the inclusion of *P. seemannii* and *P. schottii* within *P. macrocarpa* (Bernal and Galeano, 2010), instead they support the existence of six *Phytelephas* species (Barfod et al., 2010; Trénel et al., 2007).

The close phylogenetic relationship observed between *P. aequatorialis* and *P. tumacana*, and between *P. seemannii* and *P. schottii* (Figures 1, 2), has been previously inferred (Barfod et al., 2010; Trénel et al., 2007). However, the position of the Amazonian species *P. macrocarpa* and *P. tenuicaulis* is different in our analyses, resolving as sister to the other four *Phytelephas* species from the Chocó/Magdalena (Figure 2). These two Amazonian species have not been previously inferred as sister taxa, but to be related with the species from the Chocó/Magdalena (Barfod et al., 2010; Trénel et al., 2007). Therefore, two trans-Andean clades were inferred within *Phytelephas*: one in the Chocó/Magdalena and one in the Amazonia (Figure 2).

Our biogeographic analyses suggest that the ancestor of the vegetable ivory palms presented a widespread distribution in northwestern South America during the Miocene at 19.25 Ma (95% HPD 16.96–20.58; Figure 2), which agrees with a previous ancestral range estimation of Phytelepheae (Barfod et al., 2010). The Andes were not fully formed at that time and the tropical rain forests of the Chocó, Magdalena, and Amazonia were potentially connected (Hoorn et al., 2010). Early diversification in the tribe maintained the widespread distribution for the ancestor of both *Phytelephas* and *Ammandra/Aphandra* (Figure 2), as it was shown by Barfod et al. (2010). Trans-Andean diversification in *Phytelephas* at 5.27 Ma (95% HPD 5.2–5.43; Figure 2) occurred after the western Andes became a barrier between the Chocó and the Amazonia at ~10 Ma (Gregory-Wodzicki, 2000), suggesting that the western Andean uplift promoted basal diversification by vicariance in *Phytelephas*. Intraspecific trans-Andean diversification in *A. decasperma* at 7.61 Ma (95% HPD 6.42–8.61; Figure 1) can also be explained by western Andean uplift. Trans-Andean vicariance after the western Andes uplifted has been reported for several Neotropical plant and animal taxa (Carneiro et al., 2018; Luebert and Weigend, 2014; Richardson et al., 2015; Weir and Price, 2011; Winterton et al., 2014).

A second event of Andean vicariance occurred in *Phytelephas* between the Chocó and Magdalena at 4.35 Ma (95% HPD 4.28–4.84; Figure 2) although these regions have remained connected in northwestern Colombia (Hoorn et al., 2010). The diversification between the Chocó and Magdalena took place after the eastern Andes started to uplift rapidly at ~5 Ma and may have become a barrier between Magdalena and the Amazonia (Gregory-Wodzicki, 2000). Thus, the eastern Andes is not related with *Phytelephas* diversification but may have limited the colonization of the Amazonia by *P. schottii*. Further diversification in *Phytelephas* could have occurred because of adaptive shifts in pollination mechanisms and vegetative morphology (Barfod et al., 2010), but not because of Pleistocene climatic fluctuations given that all six species were already formed before 2.6 Mya (Figure 2). Three events of Andean vicariance were previously inferred to explain diversification in *Phytelephas* (Barfod et al., 2010); nevertheless, our model based in two events seems simpler and potentially more plausible.

The basal phylogenetic relationships and time estimates of the vegetable ivory palms using 32 nuclear genes were similar to those previously obtained based on few genetic markers (Barfod et al., 2010; Trénel et al., 2007). However, different and more plausible phylogenetic relationships were recovered at species level for this study, showing the utility of targeted sequence capture for inferring deep phylogenetic relationships in palms (Helmstetter et al., 2020b; Heyduk et al., 2016; Loiseau et al., 2019). Our results obtained using genome-wide markers suggest that phylogenetic relationships and estimated divergence ages between genera could be similar to those obtained using few genetic markers; nevertheless, species relationships inferred using genomic data may be different, impacting age estimates. Different phylogenetic relationships could also impact the biogeographic conclusions drawn on these trees. Fortunately, target sequence capture approaches and baits are constantly developed at lower costs (Couvreur et al., 2019; de La Harpe et al., 2019; Johnson et al., 2019), which will allow to infer more accurate phylogenetic relationships between plants species in the future.

## Acknowledgements

We thank Ecuador’s Ministerio del Ambiente for providing the sampling permit to undertake this research (MAE-DNB-CM-2018-0082). Special thanks go to Wolf Eiserhardt, Rodrigo Bernal, and María José Sanín for providing sequence data, leaf material, or occurrence records, and to Scott Jarvie for downloading occurrence records. This research was financially supported by the SENESCYT (PhD scholarship to SE), the International Palm Society (field grant to SE), the Independent Research Fund Denmark (9040-00136B to HB), the Agence Nationale de la Recherche (ANR-15-CE02-0002-01 to TLPC), and the Laboratory Mixed International BIO INCA.

## Data accesibility

The fastq sequences (R1 and R2) for all individuals have been submitted to Genbank SRA. Bioinformatic scripts used for this study can be found at https://github.com/ajhelmstetter/afrodyn.

