## Supporting information for "Phylogenomic relationships and historical biogeography in the South American vegetable ivory palms (Phytelepheae)"

**Article type:** Short communication

**Correspondence**

**SUPPORTING TABLES**

**Table S1.** Dispersal matrices used to estimate the ancestral range and biogeographic history of the vegetable ivory palms (Phytelepheae) in BioGeoBEARS (Matzke, 2013) based on the history of Andean uplift (Gregory-Wodzicki, 2000; Hoorn et al., 2010). C: Chocó, M: inter-Andean valley of the Magdalena River, A: Amazonia, O: Andes (*Ceroxylon*–outgroup).

| **0-5 Mya** | **C** | **M** | **A** | **O** |
| --- | --- | --- | --- | --- |
| **C** | 1 | 0.5 | 0.1 | 0.01 |
| **M** | 0.5 | 1 | 0.1 | 0.01 |
| **A** | 0.1 | 0.1 | 1 | 0.01 |
| **O** | 0.01 | 0.01 | 0.01 | 1 |
| **5-10 Mya** | **C** | **M** | **A** | **O** |
| **C** | 1 | 0.5 | 0.1 | 0.01 |
| **M** | 0.5 | 1 | 1 | 0.01 |
| **A** | 0.1 | 1 | 1 | 0.01 |
| **O** | 0.01 | 0.01 | 0.01 | 1 |
| **70-10 Mya** | **C** | **M** | **A** | **O** |
| **C** | 1 | 0.5 | 1 | 0.01 |
| **M** | 0.5 | 1 | 1 | 0.01 |
| **A** | 1 | 1 | 1 | 0.01 |
| **O** | 0.01 | 0.01 | 0.01 | 1 |

**Table S2.** Summary of the12 models of range evolution used to estimate the ancestral range and biogeographic history of the vegetable ivory palms (Phytelepheae) in BioGeoBEARS (Matzke, 2013). The best model (DEC) was selected based on the lowest Akaike information criterion (AIC) and AIC corrected for small sample sizes (AICc). Models are arranged from the lowest AICc to the highest.

| **Model** | **LnL** | **# params** | **d** | **e** | **j** | **AIC** | **AIC wt** | **AICc** | **AICc wt** |
| --- | --- | --- | --- | --- | --- | --- | --- | --- | --- |
| DEC | -11.24 | 2 | 0.05 | 1.00E-12 | 0.00 | 26.48 | 0.36 | 27.98 | 0.47 |
| DIVALIKE | -11.60 | 2 | 0.07 | 1.00E-12 | 0.00 | 27.19 | 0.25 | 28.69 | 0.33 |
| DEC+J | -10.87 | 3 | 0.03 | 1.00E-12 | 0.21 | 27.73 | 0.19 | 31.16 | 0.10 |
| DIVALIKE+J | -10.87 | 3 | 0.03 | 1.00E-12 | 0.21 | 27.73 | 0.19 | 31.16 | 0.10 |
| DEC null | -16.16 | 2 | 0.01 | 2.29E-03 | 0.00 | 36.31 | 0.21 | 37.81 | 0.39 |
| DEC+J null | -14.48 | 3 | 0.003 | 1.00E-12 | 0.07 | 34.97 | 0.42 | 38.40 | 0.29 |
| BAYAREALIKE+J | -14.78 | 3 | 0.02 | 1.38E-02 | 0.36 | 35.56 | 0.00 | 38.99 | 0.00 |
| DIVALIKE+J null | -14.91 | 3 | 0.01 | 1.00E-12 | 0.05 | 35.81 | 0.28 | 39.24 | 0.19 |
| DIVALIKE null | -17.52 | 2 | 0.01 | 4.88E-03 | 0.00 | 39.04 | 0.05 | 40.54 | 0.10 |
| BAYAREALIKE+J null | -16.97 | 3 | 0.01 | 1.28E-02 | 0.07 | 39.93 | 0.04 | 43.36 | 0.02 |
| BAYAREALIKE | -20.95 | 2 | 0.08 | 3.95E-02 | 0.00 | 45.90 | 0.00 | 47.40 | 0.00 |
| BAYAREALIKE null | -26.33 | 2 | 0.01 | 1.00E-02 | 0.00 | 56.67 | 0.00 | 58.17 | 0.00 |

**SUPPORTING FIGURES**

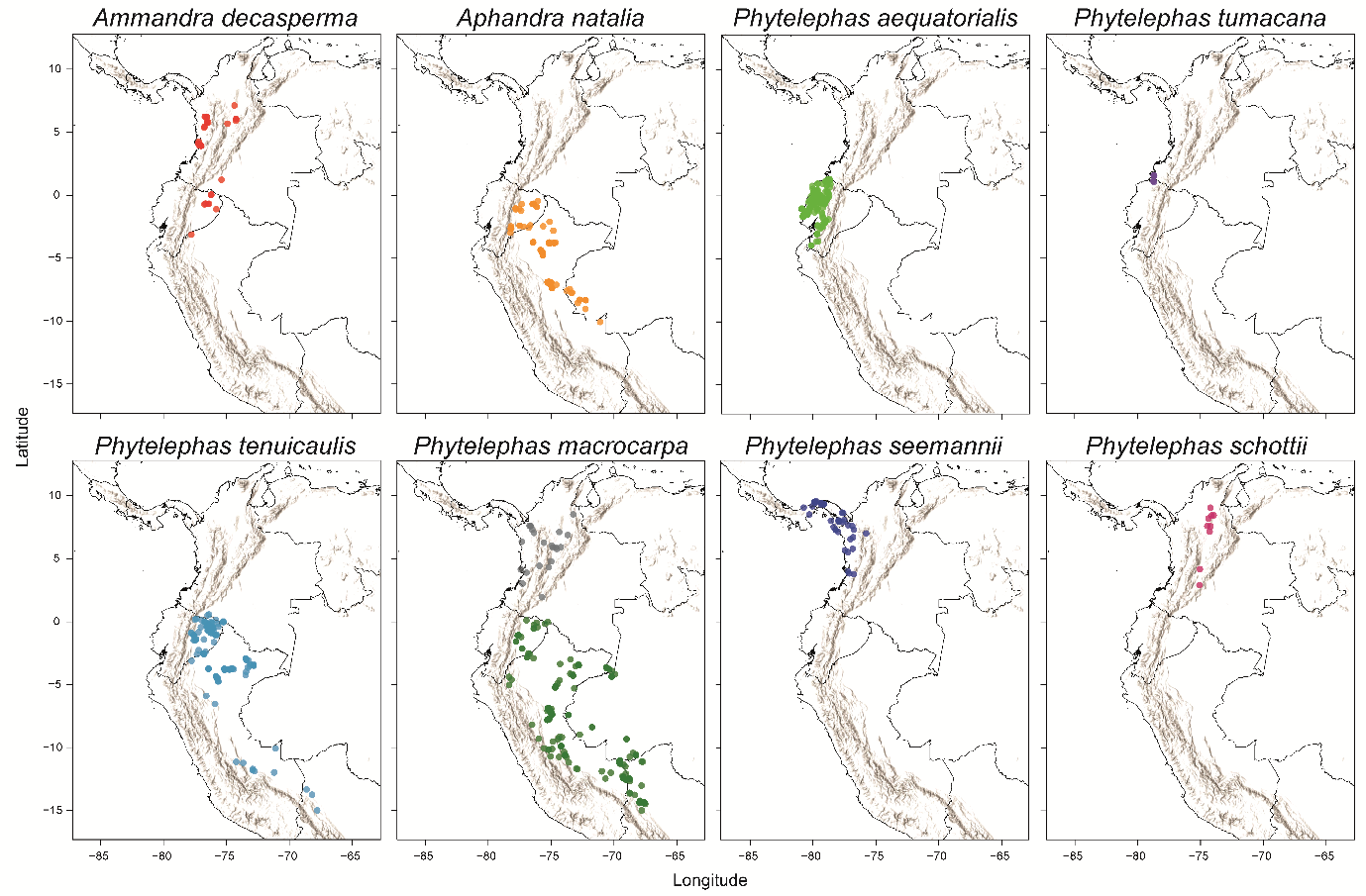

**Figure S1.** Geographic distribution of the vegetable ivory palms (Phytelepheae) in northwestern South America and adjacent Panama. Occurrence records were obtained from free online repositories and own databases, and cleaned based on specialized literature. Grey occurrence records in *Phytelephas macrocarpa* were reported in the Chocó and in the inter-Andean valley of the Magdalena River; however; our phylogenetic analyses do not support the inclusion of individuals from the Chocó within *P. macrocarpa* (Figure 1).
